# Mapping chromatin remodelling in glioblastoma identifies epigenetic regulation of key molecular pathways and novel druggable targets

**DOI:** 10.1101/2024.02.24.581853

**Authors:** Claire Vinel, James Boot, Weiwei Jin, Nicola Pomella, Charles Mein, Nicolae Radu Zabet, Silvia Marino

## Abstract

Analysis of chromatin remodelling in neoplastic stem cells as compared to ontogenetically related neural stem cells, reveals multifactorial epigenetic regulation of signalling pathways known to contribute to glioblastoma development. It also identifies novel epigenetically regulated druggable target genes on a patient-specific level, including SMOX and GABBR2 which could be further developed for future translational approaches to more effectively treat this neoplasm.

## Introduction

Glioblastoma is the most common and aggressive malignant brain tumour in the adult population. It is incurable, and patients’ survival after diagnosis rarely exceeds 15 months with relapses occurring in all patients. Despite significant research efforts, specifically in characterising the genomic, epigenomic, and molecular factors driving its development and recurrence, standard therapy including maximal safe surgical resection, radiotherapy, and chemotherapy has not changed in almost two decades.

The heterogeneous nature of the tumour, both intra-tumoural and inter-patient, is among the main therapeutic challenges of the disease. At intra-tumoural level, a subpopulation of cells called glioblastoma stem/initiating cells (GIC) has been identified from their ability to self-renew and give rise to a fully differentiated tumour upon xenotransplantation^1^. GIC significantly contribute to resistance to radiotherapy and chemotherapy ^2^, hence playing a crucial role in recurrence ^3^. Compelling evidence support an origin of GIC from neural stem cells (NSC), self-renewing and multipotent cells driving brain development and homeostasis ^1^, albeit also displaying distinct cellular and molecular features that contribute to tumour growth. At inter-patient level, distinct genetic changes characterise the tumours occurring in different patients and various subclassification of glioblastoma have been described ^4^, which however have had only a minor impact on identifying new therapies and related biomarkers so far. The contribution of epigenetic deregulation to inter-patient heterogeneity has been demonstrated, including differential methylation at the promoter of the MGMT gene which is to date the only biomarker predicting drug response in glioblastoma patients ^5^. However, despite significant efforts in characterising DNA methylation ^6^ ^7^ ^8^ and microRNA ^9^ profiles in glioblastoma, chromatin remodelling and histone modifications remain less explored, hence we still lack a global understanding of the regulatory mechanisms controlling gene expression programs in this tumour.

Modifications of histones, particularly the N-terminal tails of histone 3 (H3) at lysin residues, influence chromatin accessibility and gene expression ^11^. Acetylation of lysine 27 (H3K27ac), tri and monomethylation of lysine 4 (H3K4me3 and H3K4me1) and lysine 79 (H3K79me3), as well as trimethylation of lysine 36 (H3K36me3) are hallmarks of active chromatin, whilst methylation of lysine 9 (H3K9me3) and 27 (H3K27me3) are found at condensed/silent chromatin regions ^13^. H3K27ac and H3K4me1 marks have been linked to functional enhancers in different cell types ^14^ ^15^, whilst regions with co-localisation of H3K4me3 and H3K27me3, known as “bivalent domains”, are found mostly in embryonic stem cells ^16^. Mapping H3K27ac deposition in glioblastoma cell lines and tissue biopsies as well as in normal brain tissue revealed transcriptionally active chromatin with implications for core oncogenic dependency on super-enhancer-driven transcription factors and long noncoding RNAs ^17^. Upregulation of the histone demethylase KDM6 in GIC led to redistribution of H3K27me3 and induction of treatment resistance through acquisition of a slow cycling state ^18^. Maintenance of an undifferentiated state was shown to be achieved via bivalent modulation at lineage-specific genes ^19^ ^20^ in a highly interconnected network regulated by WNT, SHH, and HOX developmental pathways ^20^. Analysis of chromatin accessibility, histone modifications (H3K4me3, H3K27ac and H3K27me3), DNA methylation and gene expression revealed a regulatory connection between FOXM1 and ANXA2R in gliomagenesis ^21^. Finally, analysis of the distribution of H3K9ac and H3K9me3 in GIC as compared to differentiated tumour cells showed that GIC displayed an open and highly dynamic chromatin structure with loss of clustered H3K9me3 and concomitant aberrant H3K9 hyperacetylation at promoters linked to DNA damage response (DDR), thus demonstrating that the H3K9me3–H3K9ac equilibrium is crucial for GIC viability ^22^.

Here, we have leveraged SYNGN, an experimental pipeline enabling the syngeneic comparison of GIC and Expanded Potential Stem Cell (EPSC)-derived NSC (iNSC) ^23^ to identify regulatory features driven by chromatin remodelling specifically in glioblastoma stem cells. We show multifactorial epigenetic regulation of the expression of genes and related signalling pathways known to contribute to glioblastoma development. We also identify novel epigenetically regulated druggable target genes on a patient-specific level, which could be further developed for future translational approaches to tackle this neoplasm.

## Results

### Differential analysis of histone modifications in neoplastic and normal stem cells reveals epigenetic regulation of genes and molecular pathways involved in glioblastoma pathogenesis

To interrogate the global epigenetic landscape across the SYNGN cohort of 10 paired GIC/iNSC lines ^23^, we performed genome-wide chromatin immunoprecipitation sequencing (ChIP Seq) for activating and repressing histone modifications (HM), including H3K4me3, H3K27ac, H3K36me3 and H3K27me3 (Fig.S1a). H3K4me3 is predominantly enriched at promoters and transcriptional start sites (TSS) of expressed genes, H3K27ac is enriched at typical enhancer and super-enhancer regions, H3K36me3 is an elongation marker enriched in gene bodies and the PcG-catalysed H3K27me3 is involved in silencing gene expression ^24^. GIC datasets showing low number of peaks (2 tracks: GIC54 H3K36me3 and GIC61 H3K27me3, Fig.S1b) or displaying a profile clustering apart from other similar HM tracks (1 track: GIC19 H3K27ac Fig.S1c) were excluded from further analysis. Hierarchical clustering of all retained tracks shows distinct clusters in both iNSC (Fig.S2a) and GIC (Fig.S2b) for H3K27me3 and H3K36me3, but not for H3K4me3 and H3K27ac tracks in keeping with the expected overlapping distribution of these HM across the genome ^25^. Importantly, principal component analysis (PCA) of ChIP Seq data of GIC and iNSC showed a clear separation between GIC and iNSC for each HM (Fig.S3a) and correlation heatmaps of differentially bound sites highlighted differences between GIC and iNSC (Fig.S3b).

To ensure we focus on fundamental differences between the normal and neoplastic chromatin context in glioblastoma, we identified common epigenetic features shared between all 10 patients (Fig.1a). Analysis of the number of peaks of each HM (SHM analysis) showed quantitative differences with a higher proportion of H3K27ac peaks (33.3% vs 26.7%) and a smaller proportion of H3K4me3 peaks (10.3% vs 14.8%) in GIC as compared to iNSC (Fig.1b). Analysis of sites differentially enriched in each HM indicated differences between GIC and iNSC for all HM (Fig. 1c). Interestingly, genomic region annotation of the ChIP Seq peaks for each HM revealed also qualitative differences between GIC and iNSC most strikingly in H3K4me3 enrichment at promoter and 5’ UTR regions and in H3K27me3 depletion at promoter regions in GIC as compared to iNSC (Fig.1d). We then focused on the genomic regions with an expected functional impact for each HM: promoters for H3K4me3; intron and exon for H3K36me3; promoters and intron/exon for H3K27me3 and H3K27ac. Comparative analysis of the genes identified in these genomic regions in GIC and iNSC revealed that only 22 and 35% were common for H3K27ac and H3K27me3 respectively, indicating an important chromatin remodelling in GIC, which was more pronounced for these HM given that significantly more genes identified in regions with H3K4me3 and H3K36me3 peaks (43 and 45%) were common between iNSC and GIC (Fig.1e).

**Figure 1:**
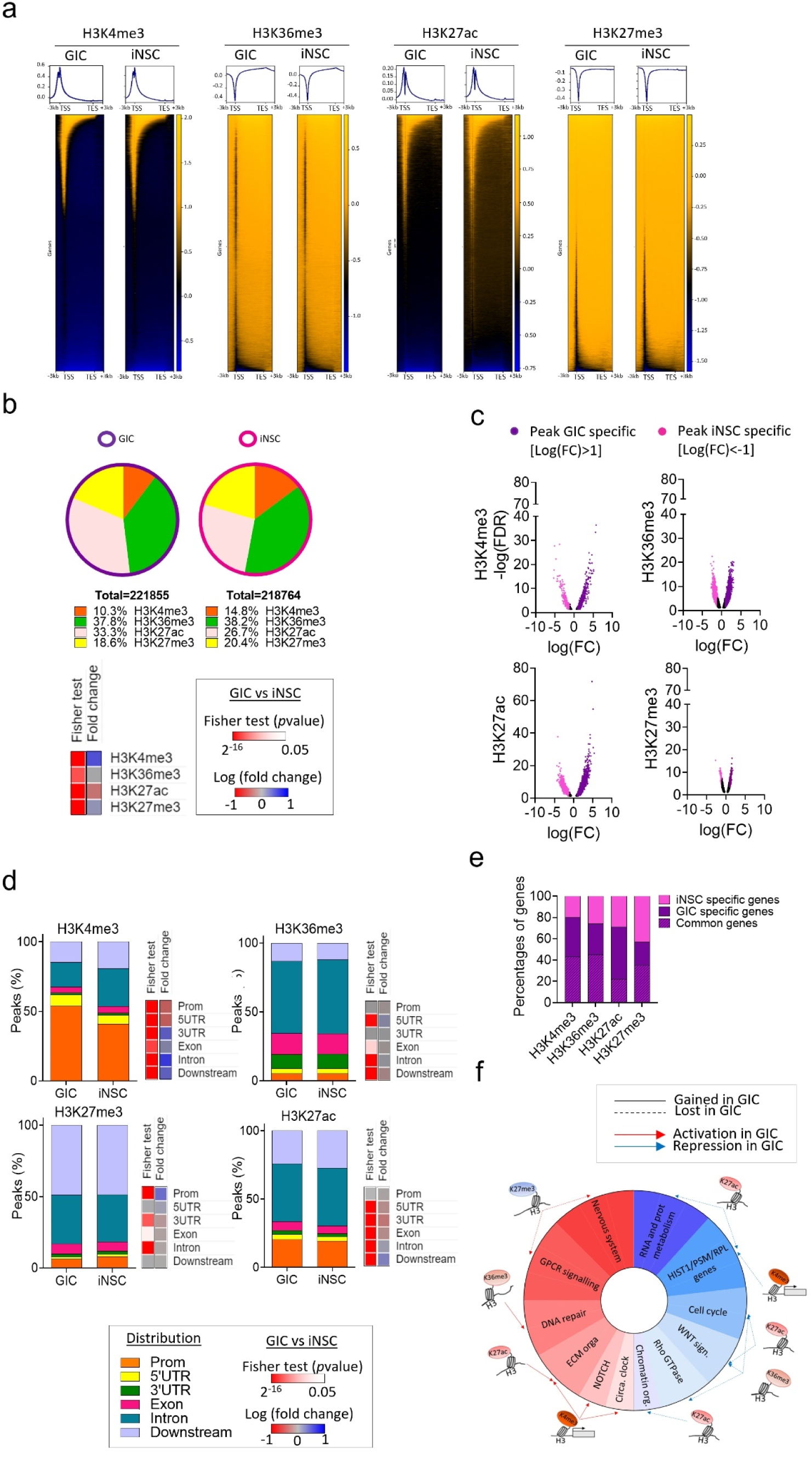
Mapping activating and repressing histone modifications in human GIC as compared to syngeneic iNSC. a) Average heat map of ChIP Seq dataset around Transcription Start Sites (TSSs) and annotated genes for the 10 patients. b) Pie charts show proportion of peaks linked to each histone modification (HM) in GIC (left) and iNSC (right) in all patients. Heatmaps represent Fisher’s exact test statistical analysis and fold change of GIC ChIP-peaks upon iNSC ChIP-peaks for each HM. c) Volcano plots represent the comparative analysis of differentially bound sites between GIC and iNSC for the four HM in all patients. Only significantly differentially bound sites are shown (FDR<0.05). Results are represented as differential log fold change of GIC upon iNSC [Log2(GIC)-Log2(iNSC). Bound sites only found in GIC are shown as purple dots (log(FC)>1) and bound sites found only in iNSC are shown as pink dots (log(FC)M<-1). Bound sites with a -1<log(FC)<1 are shown as black dots. d) Distribution of ChIP Seq peaks’ genomic annotations for each HM GIC and iNSC. Heatmap represent Fisher’s exact test statistical analysis and fold change of ChIP Seq peaks between GIC and iNSC. e) Percentages of genes common, only found in GIC (GIC specific) and only found in iNSC (iNSC specific) in each HM f) Schematic representation of HM redistribution in GIC as compared to iNSC and their targeted pathways. Red and blue shades in the donut represent activated and repressed signalling pathways, respectively.

Gene set enrichment analysis (GSEA) identified significantly enriched pathways (FDR<0.05) activated or repressed in GIC (Fig.1f) which are known to be deregulated in glioblastoma, including enrichment for neuronal system, GPCR signalling, DNA repair, extra cellular matrix organisation, NOTCH signalling and circadian clock for genes with an activating HM in GIC (Fig.1f and Fig.S4a). Conversely, genes with a repressing HM in GIC were mostly involved in metabolism of RNA and protein, HIST1, PSM and RPL cluster genes, WNT, Rho GTPase signalling as well as chromatin organisation (Fig.1f and Fig S4b).

Additionally, loss of H3K4me3 at genes of the HIST1 cluster was observed in GIC, a finding never previously reported in a glioblastoma context (Fig.S4b and table S1), and of potential interest given that epigenetic down-regulation of the *HIST1* locus has been linked to better prognosis in a proportion of acute myeloid leukemia patients ^26^. We found loss of H3K27ac at genes related to Proteasome subunits type A, B C and D (*PSM),* in particular *PSMD1* and *PSMD3* (Fig.S4b and table S1), which have been shown to act as tumour suppressors and inhibit Wnt signalling ^27^ in other cancers, though not yet described in glioblastoma. Finally, loss of H3K27ac is found in genes related to cellular response to stress, supporting the notion that cancer cells may not activate cell cycle arrest and/or apoptosis as effectively as non-neoplastic cells in a stress response context.

In summary, we show redistribution of functionally critical HM across the genome in neoplastic stem cells which affects biological processes known to play a role in glioblastoma pathogenesis, raising the possibility that epigenetic remodelling contributes to their deregulation. The approach also identifies chromatin remodelling at genes/pathways not yet linked to glioblastoma.

### Integrative analysis of single histone modifications and transcriptome reveals a direct impact on gene expression and identifies activation of a gastrulation differentiation program in GIC

To begin understanding the functional impact of the chromatin remodelling identified in GIC we integrated transcriptomic data ^23^with the ChIP Seq datasets (Fig. S5a). As expected, active regulatory regions defined by H3K4me3, H3K36me3 or H3K27ac peaks are found at upregulated genes whilst H3K27me is found mostly at downregulated genes in GIC as well as in iNSC, although here less prominently (Fig.S5a-c). However, active and repressive marks are also found at downregulated and upregulated genes respectively, raising the possibility that the expression of these genes is regulated by alternative/additional mechanisms (Fig.S5a). Interestingly, when the analysis was focused on the regions differentially bound uniquely in GIC or iNSC (Fig.2a and Fig.S5d), an enrichment of genes with an expression concordant with the HM was observed (Fig.2b). Over 90% of genes displaying a gain of activating HM (H3K4me3, H3K36me3 and H3K27ac) were upregulated in GIC as compared to iNSC, whilst 80% of genes gaining H3K27me3 were downregulated (Fig.2b top), with a similar observation also made for iNSC (Fig.2b bottom), hence suggesting that the redistribution of the HM observed is likely to have a functional impact.

**Figure 2:**
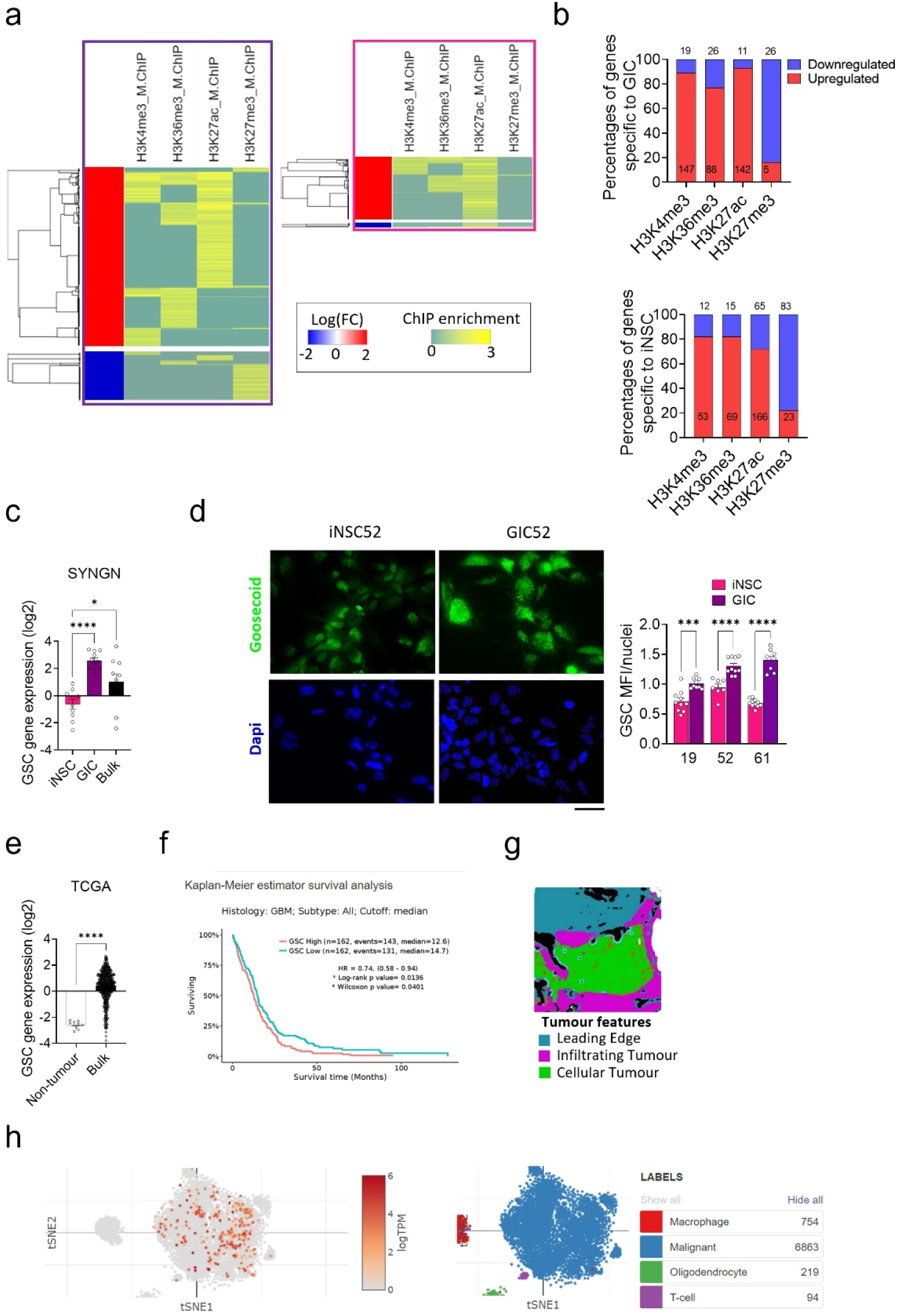
Integrative analysis of single histone modifications and gene expression reveals dynamic and synergistic epigenetic regulation of pathways involved in glioblastoma pathogenesis. a) Pearson correlation heatmaps of integrative analysis of ChIP Seq data for each HM and RNAseq dataset ^23^ for genes only found in GIC (left)or iNSC (right) for at least one HM. RNAseq data are represented as log fold change of Differentially Expressed (DE) genes between GIC and iNSC: logFC DE>1 and <-1 when genes are up (red section) and downregulated (blue section) in GIC as compared to iNSC respectively (left). LogFC DE>1 and <-1 when genes are down and upregulated in iNSC as compared to GIC respectively (right). b) Percentages of upregulated (red) and downregulated (blue) genes in iNSC as compared to GIC (top) and in GIC as compared to iNSC (bottom) for each HM based on transcriptomic dataset from the SYNGN cohort ^23^. Number of genes is also specified for each condition. c) mRNA expression of *GSC* in iNSC, GIC and bulk tumour from the RNAseq dataset of the SYNGN cohort ^23^ (left) and in bulk tumour and non-tumour samples from TCGA dataset. Results are expressed in log 2 (tpm) transcript per million (tpm). One-way ANOVA test. \**p*value<0.05, \*\**p*value<0.01 and \*\*\**p*value<0.001. d. Representative immunofluorescent images for GSC (green) in iNSC and GIC from patient 52. Nuclei are counterstained with DAPI. Scale bar: 50µm Quantification is shown as Mean Fluorescence Intensity (MFI) standardised by the number of nuclei. One-way ANOVA test. \**p*value<0.05, \*\**p*value<0.01, \*\*\**p*value<0.001, \*\*\*\**p*value<0.0001. e) GSC gene expression in non-tumour and bulk primary glioblastoma tumour (left panel). T test. \**p*value<0.05, \*\**p*value<0.01 and \*\*\**p*value<0.001. f) Survival curve of glioblastoma patients with high and low expression of *GSC* gene (right panel). Source: TCGA. Stat test: log-rank, \**p*value<0.05, \*\**p*value<0.01 and \*\*\**p*value<0.001. g) Spatial expression of *GSC* in glioblastoma bulk samples, analysed on Ivy -GAP. The left panel shows an example of histological anatomic structure identified in a sub-block and the right panel represents the expression of *GSC* in RNAseq data from anatomic structures shown as log2 normalized gene expression. Leading Edge defined as the border of the tumour, where ratio of tumour to normal cells is 1-3 / 100. Infiltrating tumour defined as the intermediate zone between leading edge and cellular tumour, where ratio of tumour to normal cells is 10-20 /100. Cellular tumour defined as tumour core, where tumour to normal cells is 100-500 / 1. One-way ANOVA test. \**p*value<0.05, \*\**p*value<0.01 and \*\*\**p*value<0.001. h) Single cell RNAseq data from *Neftel et al.* showing *GSC* expression (left panel) in scRNAseq of glioblastoma samples in clusters defined in ^60^ (right panel). Data are plotted as tSNE, with logTPM expression ranging from light orange to dark.

To identify pathways directly regulated by HM in GIC, we selected the concordant genes (activating HM/upregulation of expression and repressive HM/downregulation of expression) for each HM and performed GSEA (Fig.S5e and b, table S2). In the first instance we confirmed the results of the SHM analysis that pathways known to play a role in glioblastoma pathogenesis are epigenetically regulated, including pathways related to neoplastic transformation (Receptor Tyrosin Kinase (RTK) signalling (EBRR2) ^28^, WNT signalling ^29^ GPCR signalling ^30^), neuronal systems (potassium channels ^31^), cellular response to stress and ALK signalling ^32^ ^33^, as well as genes involved in circadian clock regulation ^34^ ^35^ ^36^ (Fig.S5e and table S2). Pathways involved in modulation of the inflammatory microenvironment via interferon signalling (antigen presentation, cytokine signalling and interferon signalling) ^37^ were also identified, in keeping with an intrinsic regulation of the inflammasome mediated by the redistribution of HM in tumour cells (Fig.S5e and table S2). Interestingly, we found decreased expression of a group of 37 ribosomal protein coding genes associated with the combined loss of the three activating histone modifications (Fig.S6a and table S2), in keeping with chromatin remodelling impacting ribosome biogenesis^38^.

Importantly, our analysis also identified novel pathways/genes specifically deregulated in GIC as compared to iNSC including pathways associated with activation of GABA B receptors signalling (GABA B receptor activation and activation of GABA B receptor) and activation of the gastrulation pathway (table S2) mediated by the gain of H3K4me3 and upregulation of Goosecoid (GSC) (Fig.S6b). We confirmed *GSC* upregulation in GIC as well as in the bulk tumour as compared to iNSC by qPCR (Fig.2c) and at protein level (Fig.2d and S6c and d). Leveraging the Cancer Genome Atlas Program (TCGA) we confirmed its upregulation in additional glioblastoma samples as compared to non-tumour tissue (Fig.2e) and importantly demonstrated a link to poorer prognosis in higher expressors in this cohort of patients (Fig.2f). At regional level (Glioblastoma Atlas Project - IvyGap), *GSC* expression is highest in the cellular tumour (tumour core) and show intermediate expression in the infiltrating tumour (ratio of tumour-to-normal cells 10-20/100) as compared to leading edge (ratio of tumour-to-normal cells 1-3/100 (Fig.2g), with *GSC* being exclusively expressed in malignant cells at single-cell transcriptomic level (Single Cell Portal, The Broad Institute) (Fig.2h).

In summary, we show that the redistribution of key regulatory HM in the neoplastic stem cell context has an impact on gene expression and identify novel epigenetic regulatory programs, including reactivation of a gastrulation differentiation program in glioblastoma.

### Chromatin states segmentation confirms the contribution of chromatin remodelling to regulation of mechanisms of glioblastoma pathogenesis

Next, we set out to capture the complexity of the epigenetic deregulation in glioblastoma in a systematic manner. Using ChromHMM genomic segmentation ^39^, we explored the unsupervised combinatorial patterns of the four histone marks in an 8-state model and predicted active and inactive chromatin states at specific genomic features in GIC as compared to iNSC (Fig.3a). The 8 chromatin states were defined as active states – “transcription”, “active transcription”, “enhancers” and “active TSS”- as well as inactive states – “quiescent”, “weak repressed polycomb”, “repressed polycomb” and “poised gene body”. We first trained the model on iNSC and applied it to GIC and vice versa (Fig.S7a) and given that a similar level of generalisation was observed (Fig.S7a), we selected the iNSC model for the downstream analysis. Assessment of percentages of the genome in each state showed that the majority was in weak transcription/quiescent state (72 and 70.3%) and 1-8% in an enhancer state (Fig.3b and S7b) in both GIC and iNSC, similar to previous work ^40^. Noticeably though, the distribution of the peaks across the states is different in iNSC and GIC with a higher and lower respective percentage of peaks in poised gene body state (1.54 vs 0.68%) and in repressed polycomb state (1.39 vs 3.01%) in GIC as compared to iNSC (Fig.3b). In the activating states, we find more peaks in the transcription state in GIC than in iNSC (5.96 vs 4.93%) whilst fewer peaks are to be found in the enhancer state (1.77 vs 2.49%) (Fig.3b), indicating a loss of enhancer activity in GIC. The genomic region annotation of the peaks for each state did not reveal striking qualitative differences between GIC and iNSC (Fig.S7c) and the comparison of peaks in each chromatin state revealed that regions are common between GIC and iNSC highlighting the similarities between the two cell types (Fig.S7d).

**Figure 3:**
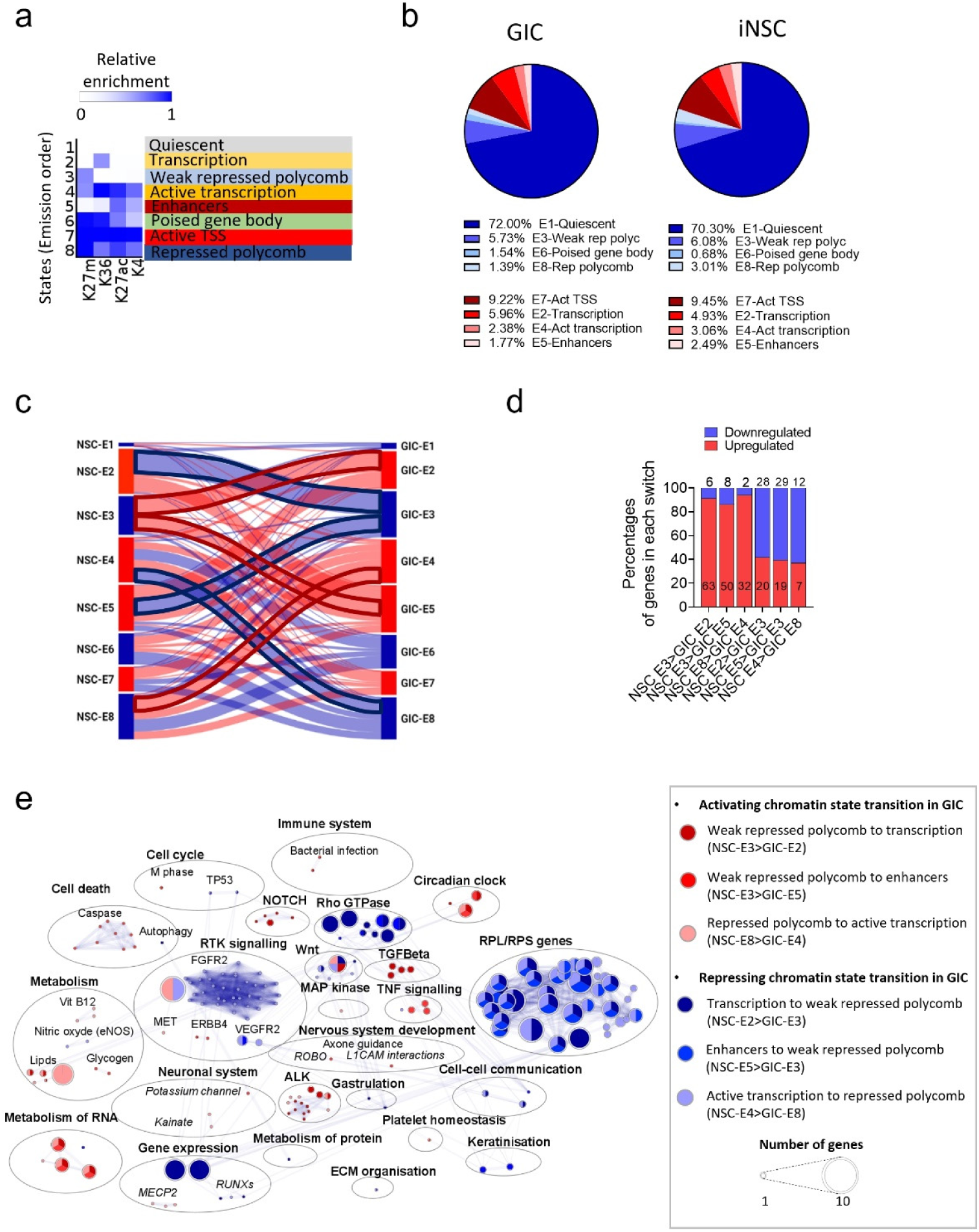
Comparative analysis of the functional impact of chromatin states dynamics in GIC and iNSC using automatic fragmentation analysis. a) Chromatin states defined by enrichment of HM using ChromHMM ^39^. Probabilities of each HM in chromatin states is depicted as a heatmap. b) Pie charts show percentages of peaks in each chromatin state in GIC (left) and iNSC (right). c) Sankey diagram shows the switch of peaks from one chromatin state in iNSC to another in GIC. The thickness of the links is proportional to the number of peaks included. Flows with the highest number of peaks between two opposites state functions are highlighted in bold red (activating transition in GIC) and blue (repressing transition in GIC). d) Percentages of upregulated (red) and downregulated (blue) genes in the chromatin states of interest based on transcriptomic dataset from the SYNGN Cohort ^23^. Number of genes is also specified for each condition. e) Visualisation of the enriched pathways identified in GIC from genes activated in GIC as compared to iNSC and from genes inactivated in GIC as compared to iNSC. Pathways are annotated based on pathways enrichment analysis performed with Reactome and represented as circle, colours represent each histone (see legend), size of the circle is proportional to the number of genes involved in the pathway (FDR<0.05).

As for the SHM analysis, we focused on the regions uniquely found in GIC and iNSC (Fig.S7d) and integrated these results with the transcriptomic data to focus on functionally relevant events. We show 87, 85, 78 and 63% of concordance between active chromatin states (respectively transcription, active transcription, enhancers and active TSS) and gene upregulation both in GIC and iNSC (Fig.S8a left) thus confirming a strong epigenetic regulation of gene expression in this cell type. However, lower percentages of downregulated genes, 63, 75 and 59%, are found in repressing chromatin states (respectively quiescent, weak repressed polycomb, poised gene body and repressed polycomb) (Fig.S7a right) in both cell types, in keeping with the more nuanced impact on transcription regulation of these chromatin states, which may lead to low reduction in gene expression which may not have been captured by the thresholds we used for DE analysis.

GSEA of concordant genes confirmed epigenetic regulation of pathways known to be contributing to glioblastoma pathogenesis such as transcription factors *RUNX* ^41^, *SMAD*s ^42^ and PPARA ^43^, neddylation of protein ^44^, homeostasis disorder including Basigin ^45^ and Kallikrein/kinin complex ^46^. Conversely, loss of repressing states in GIC leading to upregulation of genes involved in extra-cellular matrix organisation pathways (Laminin interactions, integrin cell surface interactions and ECM proteoglycans) and necrosis was observed (Fig.S8b). Enrichment of inflammasome pathways (interferon signalling, antigen presentation and cytokines signalling) and circadian clock-related pathways with gain of activating chromatin states and loss of repressing states respectively confirmed our previous finding (Fig.S8b and table S3). Within the inflammasome, the interferon-related pathways are consistently enriched because of the upregulation of two of the 2’-5’-oligoadenylate synthases family (*OAS1* and *OAS3*). *OAS1* is known to be upregulated ^47^ and hypomethylated ^48^ in glioblastoma and its silencing leads to an increase of temozolomide-sensitivity in vitro ^47^. Interestingly, we find an upregulation of *OAS1* and *3* in our dataset in GIC as compared to iNSC (Fig.S8c), that is linked to worse prognosis for *OAS3*, as indicated by poorer patient survival (TCGA database) (Fig.S8d). Regional expression dataset (IvyGap) revealed no difference between the locations analysed (leading edge, infiltrating tumour, and cellular tumour) for OAS1 with the expression not being tumour cells-specific, whilst OAS3 is significantly more expressed in the cellular tumour (tumour core) (Fig.S8e). Non-exclusive expression in the tumour cells, with immune cells including macrophages and T cell and oligodendrocytes also expressing these genes was confirmed at single cell transcriptomic level (Fig.S8f) in keeping with their immune regulatory role.

Similar to our findings in the SHM analysis, epigenetic-mediated downregulation of *RPL* and *RPS* genes is confirmed also with the ChromHMM approach, mostly mediated by the loss of the 4 activating chromatin states and the gain of poised gene body state (Fig.S9a and table S3). Signalling pathways related to the receptor tyrosine kinase MET ^49^ and ECM organisation ^50^ are also found enriched by genes epigenetically downregulated in GIC as compared to iNSC (Fig.S9a) as well as deregulation of keratinisation, previously described in a glioblastoma context ^51^. Importantly, we identified an enrichment for cellular response to hypoxia (Fig.S9a), potentially mediated by the epigenetic downregulation of the Ubiquitin-conjugating enzyme E2 D1 (*UBE2D1*) known to be responsible for the ubiquitination of hypoxia-inducible transcription factor HIF-1 alpha leading to its degradation. Whilst it has not been described in glioblastoma to date, our data suggests that in GIC, *UBE2D1* is in a poised state which could allow the cancer cells to respond quicker to cellular stress such as hypoxia by decreasing HIF1-apha degradation and promoting angiogenesis and hence tumour maintenance in a hypoxic environment. Interestingly, its low expression is linked to poor prognosis (Fig.S9b) and regional transcriptomic shows it to be expressed by malignant cells as well as other brain and immune cells (Fig.S9c).

In summary, unbiased genomic segmentation confirmed that chromatin states remodelling contributes to regulation of key oncogenic pathways in glioblastoma.

### Analysis of transitioning peaks from a repressing state in iNSC to an activating state in GIC and vice versa identifies GABBR2 and SMOX as novel druggable target genes involved in migration and invasion of tumour cells

To capture the potentially more functionally relevant chromatin conformation changes and to capitalise on the patient-specific comparison enabled by the SYNGN platform, we then focused on peaks that were transitioning/switching from one state in iNSC to another state in GIC in at least two patients of the cohort (Fig.3c and S10a). We observed switching in all 8 chromatin states with most switches found in state 4 (active transcription) in both iNSC and GIC and less switches in state 6 (weak transcription/quiescent) (Fig.3c and S10a) with more peaks observed in repressing switches as compared to activating ones (57% vs 43%) (Fig.S10b). Notably, the most frequent switches occurred within weak polycomb, transcription, active transcription, and enhancers (Fig.3c and S10a). Genome annotation of the peaks revealed that they are mostly found in promoter regions for the activating switches “weak polycomb to enhancers”, “repressed polycomb to active transcription” and repressing switches “transcription to weak repressed polycomb”, “enhancers to weak repressed polycomb”; whilst most peaks are in the intron regions for the activating “weak repressed polycomb to transcription” and the repressing “active transcription to repressed polycomb” switches (Fig.S10c). Notably, the integration with transcriptomic dataset consistently showed that the predicted functional impact of activating switches is higher than the repressing switches (over 90% concordance vs over 60%) (Fig.3d).

The 6 switches with more peaks transitioning between states were selected for further analysis, including 3 switches from an inactivating state in iNSC to an activating state in GIC (weak polycomb to transcription; weak polycomb to enhancers and repressed polycomb to active transcription) and 3 switches from an activating state in iNSC to an inactive state in GIC (transcription to weak repressed polycomb, enhancers to weak repressed polycomb and active transcription to repressed polycomb) (Fig.3c and S10a). GSEA analysis of the differentially regulated genes confirmed as significantly enriched (FDR<0.05) several pathways identified in the SHM analysis including RTK and WNT signalling, *RPL/RPL* genes, which are transitioning from activating chromatin states in iNSC (“transcription” or “enhancers”) into a repressing state in GIC (“(weak) repressed polycomb”) as are the circadian clock -related pathways (Fig.3e and table S4). It has been previously described that the downregulation of two key transcription factors BMAL1 or CLOCK in GIC induces cell cycle arrest and apoptosis ^33^. Heterodimer of CLOCK and BALM1 are a major transcriptional regulator of the circadian clock mechanism in mammals. Our analysis reveals an epigenetically-mediated upregulation of the Neuronal PAS domain protein 2 (*NPAS2*), which is a paralog of *CLOCK*, able to functionally substitute for it in the regulation of circadian rhythmicity ^52^.

Next, we reviewed the existing literature on the genes affected by these chromatin state transitions and identified 42/102 upregulated genes and 27/50 downregulated genes as never previously described in a glioblastoma context (table S5). We focused our attention on the activated genes as they could be potentially more easily targetable pharmacologically, among these 9 had been linked to low grade glioma, 7 are found to be specifically enriched in glioma among other cancers (TCGA), 7 are predicted to interact with an FDA-approved drug (DGidb query ^53^), one is linked to poor prognosis (TCGA) and 5 can be inhibited by a commercially available small molecule (Fig.4a).

**Figure 4:**
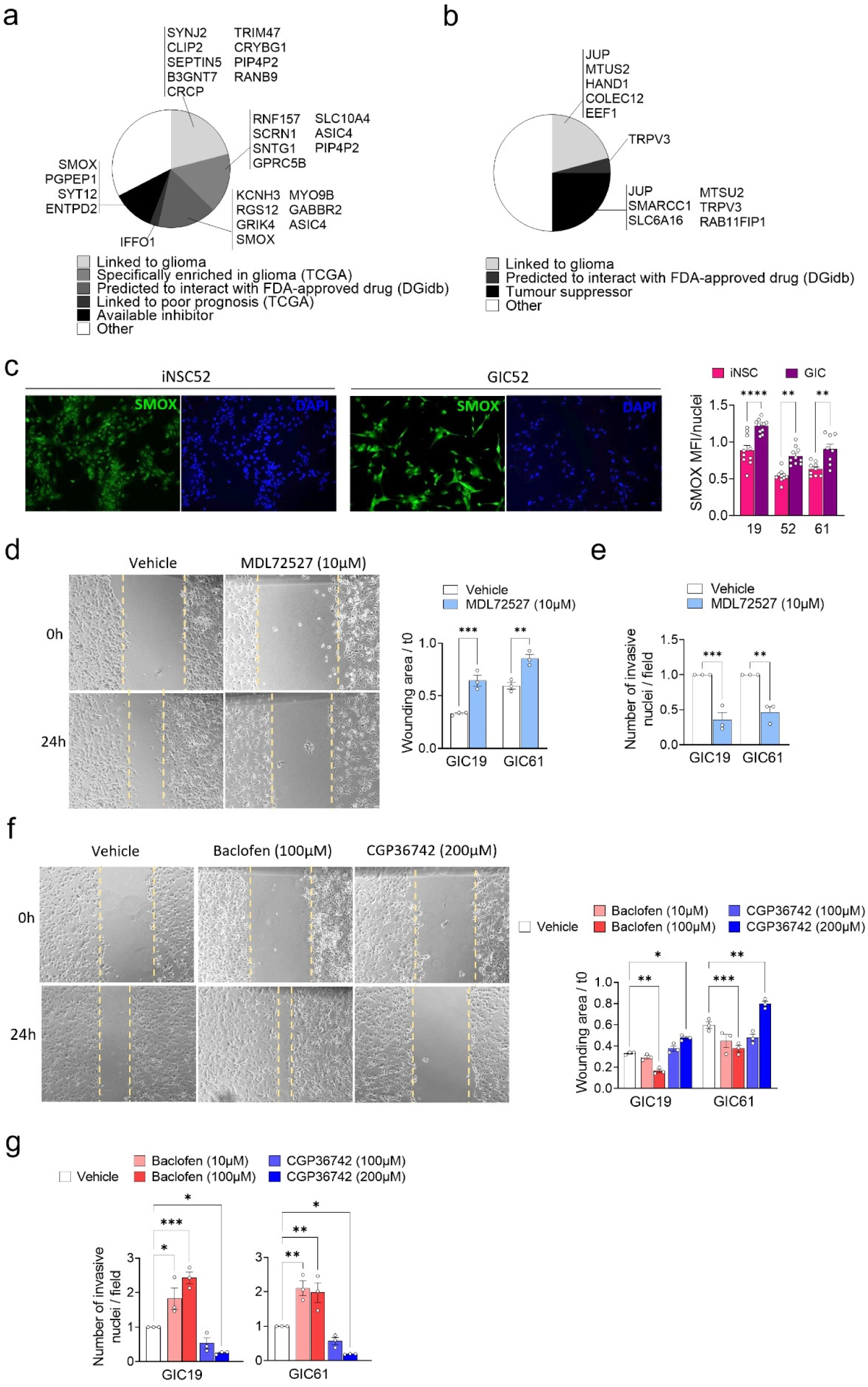
SMOX and GABBR2 are regulated by transitioning peaks in GIC. a) Venn diagrams shows the epigenetically upregulated genes newly identified in GIC classified in subgroups defined by a characteristic of interest for further molecular validation or clinical application. b) Venn diagrams shows the epigenetically downregulated genes newly identified in GIC classified in subgroups defined by a characteristic of interest for further molecular validation or clinical application. c) Representative images of SMOX protein expression (green) in iNSC and GIC from patient 52. Nuclei are counterstained with DAPI. Scale bar:100µm. Quantification of results is shown as Mean Fluorescence Intensity (MFI) standardised by the number of nuclei for three patients. One-way ANOVA test. \**p*value<0.05, \*\**p*value<0.01, \*\*\**p*value<0.001, \*\*\*\**p*value<0.0001. d) Representative bright-field images show wound healing assay in GIC19 cells treated for 24hours with 10µM of SMOX inhibitor MD72527. Quantification of results is expressed in wound area at 24 hours standardised on t0 for GIC from two patients. Experiments have been performed three times. One-way ANOVA test. \**p*value<0.05, \*\**p*value<0.01, \*\*\**p*value<0.001. e) Invasion assay quantification of results expressed in fold change of the average number of nuclei per image field of GIC from patients 19 and 61 treated with 10µM of SMOX inhibitor MDL72527 standardised on the vehicle. Experiments have been performed three times. One-way ANOVA test. \**p*value<0.05, \*\**p*value<0.01, \*\*\**p*value<0.001. f) Representative bright-field images show wound healing assay in GIC19 cells treated for 24 hours with 100µM of GABBR2 agonist Baclofen or 200 µM of GABBR2 antagonist CGP36742. Quantification of results is expressed in wound area at 24 hours standardised on t0 for GIC from two patients. Experiments have been repeated three independent times. One-way ANOVA test. \**p*value<0.05, \*\**p*value<0.01, \*\*\**p*value<0.001. g) Invasion assay quantification of results expressed in fold change of the average number of nuclei per image field of GIC from patients 19 and 61 treated with 100µM of GABBR2 agonist Baclofen or 200 µM of GABBR2 antagonist CGP36742 standardised on the vehicle. Experiments have been repeated three independent times. One-way ANOVA test. \**p*value<0.05, \*\**p*value<0.01, \*\*\**p*value<0.001.

Spermine oxidase (SMOX) catalyses the oxidation of spermine to spermidine and is found to transition from repressed polycomb to transcription through the gain of H3K27ac and H3K36me3 (Fig.S11a). Spermidine is a polyamine, known to be increased in brain tumours including glioblastoma and medulloblastoma ^54^, although the role of SMOX in this process hasn’t been characterised. Regional and single cell transcriptomic reveal that *SMOX* is highly expressed in the tumour core as well as in peripheral areas where it is mostly expressed by malignant cells (Fig.S11b and c). Gene expression analysis by qPCR (Fig.S11d) and by immunostaining (Fig.4c and S11e) confirmed higher expression of *SMOX* in GIC as compared to iNSC in three patients of the SYNGN cohort. We next aimed to modulate *SMOX* activity via a pharmacological approach with its inhibitor MDL72527 which is able to penetrate the blood brain barrier and has been shown to exert a cytotoxic effect on colon carcinoma cell lines ^55^. We show that SMOX inhibition does not affect the proliferation of GIC or iNSC (Fig.S11f), however, it promotes tumour cells migration, as assessed by wound healing assay, including assessment of gap closure (Fig.4d) and invasion (Fig.4e and S11g).

*GABBR2* has been consistently identified in all the analyses performed in our study and is found to transition from repressed polycomb to enhancers through the loss of H3K27me3 and the gain of H3K27ac (Fig.S12a). We confirmed upregulation of its expression in GIC as compared to iNSC in three patients of the cohort (Fig.S12b) and regional and single cell transcriptomic analysis revealed that it is highly expressed in the tumour core as well as in peripheral areas where it is expressed by malignant cells as well as other brain and immune cells (Fig.S12c,d). To explore a potential functional impact of GABBR2 modulation in GIC we selected an agonist (BACLOFEN, a myorelaxant antispastic agent) and an antagonist (CGP36742, selective and potent compound able to penetrate the brain). We show that pharmacological modulation of GABBR2 does not affect proliferation of tumour or non-neoplastic cells (Fig.S12e) whilst its activation promotes migration (Fig.4f top) and invasion of the tumour cells (Fig.4g top and S12f) in keeping with previous finding in breast cancer ^56^. Importantly an opposite effect is observed when GIC cells are treated with the antagonist CGP36742, with inhibition of migration (Fig.4f bottom) and invasion (Fig.4g bottom).

Comparative analysis of switching chromatin states between GIC and iNSC identified novel pharmacologically targetable genes, including *SMOX* and *GABBR2* in a subgroup of glioblastoma, which could be further explored as new therapeutic approaches to counteract glioblastoma invasiveness.

## Discussion

Taking advantage of pairs of glioblastoma stem cells and their ontogenetically linked patient-matched neural stem cells, we identified significant chromatin remodelling in glioblastoma which is primarily driven by H3K27ac and H3K27me3, given the lower number of shared genes identified in these genomic regions between normal and neoplastic stem cells, as compared to regions decorated by H3K4me3 and H3K36me3 peaks. Overall, we found a strong concordance between HM/chromatin states and gene expression and globally we found a positive correlation between activating states and mRNA levels and anticorrelation between repressive states and mRNA levels, suggesting that our profiling captures the chromatin and transcriptional state of GIC. Complementary analytical approaches of this combination of HM integrated with transcriptome analysis defined functional regions of the GIC genome with increased or reduced expression of genes uniquely in GIC. Notably, in our datasets integration with the transcriptome consistently showed that the predicted functional impact of activating states/switches is higher than the repressing ones, suggesting that the chromatin state serves as a mechanism to maintain the constitutive high expression of genes important for neural stem cell identity and function. Redistribution of the Polycomb-mediated H3K27me3 mark was also observed, consistent with a model whereby GIC identity is in part due to H3K27me3 mediated silencing of genes although other mechanisms, such as for example H3K9me3, must contribute to downregulation of gene expression in these cells, particularly in a non-neoplastic context, given the lower concordant correlation with gene expression. Our findings support and further expand previous studies which have characterised the active enhancer landscape in GIC and primary glioblastoma tissue by means of integrative analysis of the HM H3K27ac and other (epi)genetic datasets with gene expression ^57^.

It is of interest that several of the molecular processes known to be involved in glioblastoma pathogenesis are either confirmed or identified as epigenetically regulated in this study. NOTCH 1 signalling has been previously described as activated in glioblastoma, including via epigenetic regulation mediated through H3K4me3 ^58^, and our data confirm this regulation hence also lending support to our experimental approach. We show loss of H3K36me3 and concomitant repression of the beta-catenin degradation pathway via Axin in keeping with existing knowledge of over activation of WNT pathway in glioblastoma ^29^. Genes involved in tumour progression including non-integrin membrane-ECM interactions, *COL4A* and *PDGFA,* showed a gain of H3K4me3 in our dataset, in keeping with epigenetic regulation of these mechanisms being critical. GIC are equipped with a robust DNA repair mechanism mainly mediated by Homologous Recombination Repair (HRR) pathway ^59^, and we show its epigenetic regulation by gain of H3K36me3; as well as pathways involved in increased genomic instability through alteration of NHEJ/HDR pathways, which we show to be at least in part epigenetically regulated through the gain of H3K36me3. We identified an enrichment for cellular response to hypoxia, potentially mediated by the epigenetic downregulation of the Ubiquitin-conjugating enzyme E2 D1 (*UBE2D1*) known to be responsible for the ubiquitination of hypoxia-inducible transcription factor HIF-1 alpha leading to its degradation (provided by RefSeq, Mar 2011). Whilst it has not been described in glioblastoma to date, our data suggests that, in GIC, gene body region of *UBE2D1* is in a poised state which could allow the cancer cells to response quicker to cellular stress such as hypoxia by decreasing HIF1-apha degradation, promoting angiogenesis and tumour maintenance in a hypoxic environment. Interestingly, its low expression is linked to poor prognosis (Source: TCGA) and spatial expression shows its expression by malignant cells as well as other brain and immune cells ^60^. Our data suggest that the poor prognosis could be tumour-cell driven, given the epigenetic regulation observed in GIC, however a contribution of the tumour microenvirorment cannot be excluded given the spatial expression pattern observed.

Interestingly, we also identified pathways regulated by a combination of histone modifications, gain of H3K4me3, H3K36me3 and H3K27ac or loss of H3K27me3 respectively, including cellular response to stress and ALK signalling, for which a dysregulation has been described in glioblastoma ^32^ ^33^. Pathways involved in the modulation of the inflammatory microenvironment via interferon signalling (antigen presentation, cytokine signalling and interferon signalling) which were shown to regulate cell death and mesenchymal phenotype ^37^ are epigenetically regulated in GIC, hence demonstrating a cell intrinsic regulation of the inflammasome mediated by the redistribution of HM in tumour cells. In 2019, Dong et al showed that GIC can be targeted through the downregulation of the circadian clock genes *BMAL1* or *CLOCK* leading to cell-cycle arrest and apoptosis ^34^. Moreover, CLOCK has been shown to promote tumour angiogenesis ^35^ and immunosuppression ^36^. We show here that genes involved in circadian clock are epigenetically upregulated in a multi-factorial fashion by means of gain of the 3 activating HM included in our analysis (H3K4me3, H3K36me3 and H3K27ac).

Our study shows an epigenetic regulation of a large group of 57 ribosomal protein coding genes mediated by the loss of H2K27ac, 37 of which also showed decreased expression associated with the combined loss of the three activating HM. Dysregulation of ribosomal proteins (RPL/RPS) is a hallmark of cancer including glioblastoma ^38^ ^61^ and importantly ribosome biogenesis has been described to support the synthesis of protein involved in the differentiation process of NSC ^62^. Downregulation of the expression of ribosomal genes contributes to myeloid lineage differentiation in bone-marrow derived-macrophages ^63^ and a recent study showed that GIC acquire an epigenetic immune editing process launching a myeloid-affiliated transcriptional program as an immune evasion program ^64^, hence it is possible to speculate that epigenetic regulations such as those described here could contribute to this plasticity.

Developmental programs participating in tissue development and homeostasis re-emerge in tumours including glioblastoma, and reactivation of such aberrant expression programs supports stemness, growth and migratory properties of the tumour cells ^65^. We show loss of H3K27me3 and gain of H3K4me3 at the pluripotency markers, *NANOG, OCT3/4* and *SOX2*, in GIC, hence maintaining their embryonic-like gene expression signature, which contributes to drug resistance and recurrence ^66^. Interestingly, we also identified loss of H3K27me3 in genes involved in pancreatic beta-cell development (*ONECUT3, ONECUT1, NKX6-1, NKX2-2, RFX6, MAML3, PTF1A, FOXA3, NEUROG3*) raising the possibility that the redistribution of H3K37me3 in GIC could lead to reactivation of the beta-cell transcription programme in these cells with potential impact on insulin production and glucose metabolism. This supports previous findings showing that activation of insulin-mediated signalling pathways in glioblastoma promotes proliferation and survival of the tumour cells through PI3K/Akt ^67^ and anti-glycemic therapy has been recently shown to enhance PI3K inhibitor efficacy in glioblastoma patients. ^68^. Finally, we identified the H3K4me3 mark and activation of the expression of Goosecoid (GSC) in GIC and demonstrated a link to poorer prognosis in higher expressors in the TCGA cohort of patients. GSC is an homeobox gene expressed specifically in the dorsal blastopore lip of the gastrula which plays an important role in the Spemann’s organizer formation. Its upregulation promotes tumour progression in several cancers, including breast ^69^, colorectal ^70^ and hepatocellular carcinoma ^71^ by promoting invasion and migration, although the underlying mechanism is unclear. It is intriguing that the gastrulation process is characterised by dynamic cell movements when the blastula differentiates into three germ layers (endoderm, mesoderm and ectoderm) and our data raise the possibility that this mechanism is aberrantly re-activated in glioblastoma through epigenetic deregulation promoting motility to facilitate spread ^72^.

Epigenetic deregulation is an attractive albeit challenging therapeutic target for tumours, such as glioblastoma, where it is prominent. In fact, the only biomarker predicting response to a treatment which has some success in treating glioblastoma is methylation of the MGMT promoter predicting response to TMZ ^5^. Our study has identified two novel druggable target genes, *SMOX* and *GABBR2*, which are differentially regulated and expressed in normal and neoplastic stem cells in glioblastoma. We show that their pharmacological inhibition impacts cell migration and invasion exclusively in the neoplastic stem cells, and further testing aiming at assessing their suitability as novel druggable target genes in glioblastoma is warranted.

## Acknowledgements

This work is funded by grants from Brain Tumour Research (Centre of Excellence award to S.M.), Cancer Research UK (C23985/A29199 programme award to S.M.), Barts Charity (MGU0447 programme grant to S.M.) and The Brain Tumour Charity (GN-000717 Future Leaders award to C.V.). We thank Maeve McLaughlin, Blizard Advanced Light Microscopy Facility for sharing her expertise. We acknowledge the use of data generated by the TCGA Research Network: https://www.cancer.gov/tcga, and Ivy Glioblastoma Atlas Project (https://glioblastoma.alleninstitute.org/).

## Author contribution

CV designed, performed and analysed all wet lab experiments, contributed to the analysis and interpretation of the computational results, co-wrote the paper; JB analysed and interpreted all ChIP Seq and RNASeq data with contributions from WJ and NP; NRZ supervised ChIP Seq and RNA-Seq data analysis and contributed to the interpretation of the computational results; SM conceived the project, secured financial support, supervised the experiments and co-wrote the paper with contributions from all other authors.

## Data availability

**All in-house python codes are available on Github (**https://github.com/wjin722/chip_seq_analysis?tab=readme-ov-file).

## Material and methods

### Human cell culture

Human primary GIC and iNSC cells originated from a novel experimental pipeline previously generated to derive cells from patients who underwent surgical resection of glioblastoma ^23^. The use of human tissue samples was approved by the National Research Ethics Service (NRES), University College London Hospitals NRES Project ref 08/0077 (S Brandner); Amendment 1 17/10/2014. Briefly, GIC were isolated from fresh tumour tissue following a published protocol ^73^ and fibroblast from a small strips of dura mater. Fibroblasts were then reprogrammed into EPSCs ^74^, which were induced into iNSC following manufactured protocol (Gibco, #A1647801).

GIC were cultured on laminin (Sigma, #L2020) coated tissue plates in NeuroCult NS-A proliferation kit media (Stem Cell Technologies Cat. #05751) supplemented with 1% penicillin/streptomycin solution (Sigma Cat. #P4458), heparin (2µg/ml, Stem Cell Technologies Cat. #07980), mEGF (20ng/ml, Peprotech Cat. #AF-315-09-1MG) and hFGF (10ng/ml, Peprotech Cat. #AF-100-18B-50UG) and dissociated with accutase (Millipore, #SCR005) once they reached 70% confluence for replating. Cells were stored in liquid nitrogen in Stem Cell Banker freezing media (Ambsio ZENOAQ, #11890).

iNSC were cultured on geltrex (Gibco, #A1413302) coated tissue plates in Neural expansion media made of Neurobasal 0.5X (Gibco, # 21103049), Advanced™ DMEM⁄F-12 0.5X (Gibco, #12634010) supplemented with 1% penicillin/streptomycin solution and neural induction supplement (Gibco, #A1647801) and dissociated with accutase for replating. Cells were stored in liquid nitrogen in Synth-a-Freeze cryopreservation medium (Gibco, #A12542). All cells were cultured at 37°C, 5% CO.

### RT qPCR

Total RNA was extracted from the cell pellet with RNeasy kit (Qiagen 74004) following the manufacturer’s protocol. 0.5 μg were retrotranscribed by SuperScriptIII (Invitrogen, 18080093). 5 ng of cDNA template and SYBR Green primers were used to perform SYBR Green assay using SYBR Green PowerUp Master Mix (Applied Biosystems, A25742) and run on a StepOne Real-Time PCR System (ThermoFisher). The housekeeping genes GAPDH or ATP5B were used. GAPDH: FW 5’-CTGAGGCTCCCACCTTTCTC-3’; REV 5’-TTATGGGAAAGCCAGTCCCC-3’, ATP5B: FW 5’-GCGAGAAGATGACCCAGATC-3’; REV 5’-CCAGTGGTACGGCCAGAGG-3’, SMOX: FW 5’-TAACTCGTGACCTCCAGC-3’; REV 5’-GCGGCTAGCTCTACAGAA-3’, GABBR2: FW 5’-GACCATCTCAGGAAAGACTC-3’; REV 5’-GGTCTCGTTCATGGCATT-3’.

### Immunofluorescence

Adherent cultures of GIC and iNSC on a glass coverslip were washed once with PBS then fixed with PFA4% for 30 min at room temperature. After three washes with PBS, blocking with 10% goat normal serum, 0.1% Triton X100 PBS for 30 min at room temperature was performed prior to incubation with primary antibodies: 1/100 anti-SMOX (ThermoFisher, PA5-100112), anti-GSC (ThermoFisher, MA5-38019) at 4° overnight. After three washes with PBS and 1h incubation with 1/200 secondary antibodies diluted in PBS, cells were washed again with PBS, and coverslip were mounted on SUPERFROST slides with mounting media containing DAPI for nuclear counterstaining (ProLong™ Gold Antifade Mountant, Invitrogen P36930). Microscope analysis was performed with Leica DM5000 EpiFluorescence.

### Drug treatment

In vitro drug treatments in adherent cells were performed on 5k cells plated in 96 well plates coated with geltrex and laminin for iNSC and GIC culture respectively. A three days treatment was performed with increasing doses of GABBR2 agonist CGP36742 (MedChem Express HY-121599 at 100 and 200μM), GABBR2 antagonist Baclofen (Tocris, 0417 at 10 and 100 μM) and SMOX inhibitor MDL72527 (Sigma, M2949 at 10 μM) or vehicle. At end-point, cell viability and cytotoxicity were measured with CellTiter-Glo Luminescent Cell Viability Assay with CellTox Green Cytotoxicity Assay (Promega kits G7570 and G8741).

### Wound healing migration assay

5x10^4^ cells of GIC were seeded per well in a 96-well plate coated with laminin (Sigma #L2020). The next day, a vertical scratch was made with a 10μl tip and 50 µg/mL mitomycin C (Sigma Aldrich) was added to the growth medium to inhibit the cell proliferation. Picture at 10X magnification was taken following the scratch, as D0, and location of the picture was marked for each well on the plate’s lid. Pictures were taken at marked location for each well after 24hours incubation. Surface area of the wound was calculated with Image J. Experiments were performed 3 times with 2 technical replicas for each time.

### Invasion assay

Transwell inserts with 8.0µm pores (Sarstedt Cat. #89.3932.800) were placed into wells of a 24-well plate and coated with 100µL of GelTrex. A total of 100,000 GIC cells were then seeded into the transwell insert in 200µL media with additional 700µL of normal growth media was added to the bottom chamber of the well. Cells were then incubated in normal growth conditions for 24hr, at which point cells on the inside of the transwell were removed using a cotton bud dampened with DPBS. Once cells inside the transwell were removed, cells on the bottom of the transwell were fixed using methanol, pre-chilled at –20°C, for 5 min at room temperature. After fixation, the bottom of the transwell was washed twice for 5 min using DPBS. The membrane of the transwell was then cut out and mounted onto a microscopy slide with mounting media including DAPI (ProLong™ Gold Antifade Mountant, Invitrogen P36930). Transwell membranes were then analysed at the microscope and five representative images of nuclei on each membrane captured. Each experiment was repeated three times at different passages for each cell line. Each time, two technical replicate membranes were imaged. Finally, the number of nuclei in each image field was counted, using ImageJ software, to ascertain how many cells migrated across the membrane.

### RNA extraction, RT and qPCR

Total RNA was isolated from cell pellets with RNeasy Micro purification kit (Qiagen, 74104) and digested with DNaseI (Applied Biosystems). The cDNA synthesis was carried out with SuperScript III Reverse Transcriptase Kit (Invitrogen) following manufacturer’s protocol. Analysis of gene expression was performed with the Applied Biosystems 7500 Real-Time PCR System using SYBR Green PCR Master Mix (Applied Biosystems) according to standard protocols. Technical triplicates for each sample were analysed. The Ct values of all the genes analysed were normalized to the average Ct of ACTB and *ATP5F1B* and 2ΔCts were calculated on iNSC value. Primers used in SYBR Green qPCR are the following. *ACTB*: FW 5’-GCGAGAAGATGACCCAGATC-3’, REV 5’-CCAGTGGTACGGCCAGAGG-3’; *ATP5F1B*: FW 5’-CCCAGGCTGGTTCAGAGGT-3’, REV 5’-AGGGGCAGGGTCAGTCAAG-3’; *SMOX*: FW 5’-TAACTCGTGACCTCCAGC -3’, REV 5’-GCGGCTAGCTCTACAGAA -3’ and *GABBR2* FW 5’-GACCATCTCAGGAAAGACTC -3’, REV 5’-GGTCTCGTTCATGGCATT -3’.

### Statistical analysis for the wet lab experiments

Sample processing was carried out blinded. Statistical analysis was performed using GraphPad software unless otherwise stated. Significance was determined with t-test, one-way ANOVA (with Sidak’s test), or two-way ANOVA as appropriate, and displayed as the mean ± standard error (SEM). p < 0.05 was considered significant. Significance was indicated with asterisks: *p < 0.05; **p < 0.01; ***p < 0.001. Outliers were considered those data points furthest from the median value.

### Chromatin immunoprecipitation (ChIP) assays

H3K4me3, H3K36me3, H3K27ac and H3K27me3 ChIPs were performed using ChIP-IT High Sensitivity kit (Active Motif) following the manufacturer’s instructions. Briefly, 6 × 10^6^ cells were fixed with the formaldehyde-based fixing solution for 15 min at room temperature and lysed with provided lysis solution supplemented with protease inhibitors. Next, nuclei pellets were lysed, and chromatin sonicated with Bioruptor Plus sonication device (Diagenode) to obtain DNA fragments within the recommended 200–1200-bp range. In total, 25µg of sheared chromatin was then incubated with 4µg of antibody against H3K4me3 (Diagenode, C15410003-50) H3K36me3 (Abcam, ab9050), H3K27ac (Abcam, ab4729) or H3K27me3 (Diagenode C15410195) overnight at 4 °C with rotation. Following incubation with Protein G agarose beads, bound chromatin was washed, eluted and purified following the manufacturer’s protocols. Validation by qPCR-ChIP on target genes was done before proceeding to sequencing. ChIPed DNA was end-repaired, A-tailed and adapter-ligated before size selection and amplification. The obtained libraries were QC’ed and multiplexed before 75-bp paired-end sequencing on HiSeq4000 (Illumina).

### Computational analysis of ChIP Seq datasets

The quality of ChIP Seq samples was first assessed via FastQC and TrimGalore, removing low-quality and adapter sequences. The average Phred score of the surviving reads across all samples was 30 and the average sequencing depth was 36.3 M (min = 22.1 M, max = 56.7 M). The alignment to the Ensembl GRCh38 human reference genome was performed via Bowtie v2.3.4 ^75^ with default parameters, in concomitance with the usage of samtools ^76^ for the post processing and sorting of the Binary Alignment Map (bam) files. Exploratory tools such as deeptools ^77^, plotCorrelation, plotPCA and plotFingerprint on Python v2.7.15 were used to further assess sample characteristics and to address the potential presence of outliers. After performing post-alignment quality checks, peaks were called via the MACS2 algorithm ^78^ using the corresponding input background. The shifting model was disabled to make different datasets comparable and the “–broad” option was enabled for the analysis of the histone mark H3K27me3. A minimum fold enrichment of 2 was selected with an FDR of 0.05, in both narrow peak and broad peak (–broad-cut-off) statistical analyses. The Bioconductor packages in R GenomicRanges ^79^ and ChIPSeeker ^80^ were used to find regions of consensus peaks between the two cell lines, for each antibody/condition, and to annotate them based on the location with respect to the nearest transcription start site (TSS): promoters (within 3 Kb from the TSS), exons, introns, 5′ UTR, 3′ UTR and distal intergenic. The latter was merged with so-called downstream regions. The versions of all relevant Bioconductor packages were compatible with R v3.5.3.

### ChIP Seq heatmap generation

The deeptools algorithms ^81^ bamCompare, bigWigCompare and plotHeatmap were used to produce relevant bigwig files and heatmaps, to assess region-wide and genome-wide coverage. Specifically, coverage tracks were first obtained by normalising each sample alignment file against the corresponding input (--operation log2ratio) and then pooled at both replicate and cell line levels (--operation mean), to obtain a representative sample for each antibody/condition. The regions chosen to be visualised in the heatmaps were those of consensus peaks (details in the figure legends), centred on their nearest TSS.

### Differential analysis for ChIP Seq data

The differentially bound sites (DBS) comparing GIC to iNSC were found using the R package DiffBind with default parameters. Finding regions with different functionality by identifying the various combinations of histone markers (e.g. H3K27ac lacking H3K4me3 for potential enhancer) were conducted using HOMER v4.11. Finally, R packages ChIPpeakAnno and Hsapiens.UCSC.hg38 were used to annotate the functional regions. The annotated peak results were compared to the list of differentially expressed genes from the RNA-seq ^23^. A Differentially Expressed (DE)/Differentially Bound Site (DBS) pair is considered if the gene regulation in RNA expression reflects the functionality of the bound site (e.g. activated promoter leads to upregulation in RNA expression). The above analyses were done considering each patient of the cohort as a biological replica.

### Integration between ChIP Seq and RNA-seq data

The integration analysis for ChIP Seq and RNA-seq was performed using the R package Rcade. The differentially expressed (DE) RNA were obtained using R package limma, with an absolute fold change greater than 2.0 and a p-value lower than 0.01. In-house Python code (Python 3.7) was used for visualisation.

### ChromHMM fragmentation

ChromHMM (v1.18) was used to train and annotate chromatin states ^82^. Binarised input files were generated, using the BinarizeBed command, from .bed files of each of the 4 histone marks all GICs and separately all iNSCs. Binarised files were then used as input for the LearnModel command. In brief, the LearnModel command was run separately on iNSC and GIC ChIP Seq data and for a different number of states – the number of states tested for was 4, 6, 8, 10, 12, 14 and 16. Emission parameters of the trained models were compared in order to select a number of states which produced known chromatin states, for instance: quiescent, active transcription, enhancer, repressed polycomb. The number of states selected was 8. Once an appropriate number of states were selected, models trained on iNSC and GIC were compared to evaluate which model best generalised to both cell types, it was found that both models generalised similarly well, however,, the iNSC trained model was used in segmentation of samples. Using the selected model, all GIC data merged together, all NSC data merged together and each GIC and NSC line individually, were segmented and the enrichment of each chromatin state determined using the MakeSegmentation and OverlapEnrichment commands. Finally, the enrichment of each state relative to the TSS and TES were determined using the NeighborhoodEnrichment command. Segmented peaks were annotated in R (v4.2) using the ChIPSeeker package (v1.34.1) ^83^.

### ChIP Coverage Heatmaps

ChIP coverage heatmaps were generated for GIC and NSC ChIP Seq data using the deeptools (v3.5.2) package ^77^. For each ChIP marker bigwig files for each GIC sample were averaged using the bigwigAverage function. Next, signal distribution was computed using the computeMatrix and the scale-regions option. Other options specified were to set the before region start length and after region start length to 3000bp, bin size was set to 100bp, score type was set to “mean” and region body length was set to 1000.

### Gene set analysis

G profiler tool was ^84^ used to assess biological pathways terms that showed significant enrichment in the various gene sets. The enrichment for each term was deemed statistically significant if the adjusted p-value (FDR) was lower than 0.05. Cytoscape v.3.7.215 was used to visualise relevant biological networks of enriched pathways, together with EnrichmentMap application. Several layout parameters were tuned to achieve the current Cytoscape visualisation.

## References

1. Singh, S. K. et al. Identification of human brain tumour initiating cells. Nature 432, 396–401 (2004).

2. Lathia, J. D., Mack, S. C., Mulkearns-Hubert, E. E., Valentim, C. L. L. & Rich, J. N. Cancer stem cells in glioblastoma. Genes Dev. 29, 1203–1217 (2015).

3. Liu, Q. et al. Molecular properties of CD133+ glioblastoma stem cells derived from treatment-refractory recurrent brain tumors. J. Neurooncol. 94, 1–19 (2009).

4. Verhaak, R. G. W. et al. Integrated genomic analysis identifies clinically relevant subtypes of glioblastoma characterized by abnormalities in PDGFRA, IDH1, EGFR, and NF1. *Cancer Cell* **17**, 98–110 (2010).

5. Hegi, M. E. et al. MGMT gene silencing and benefit from temozolomide in glioblastoma. N. Engl. J. Med. 352, 997–1003 (2005).

6. Klughammer, J. et al. The DNA methylation landscape of glioblastoma disease progression shows extensive heterogeneity in time and space. Nat. Med. 24, 1611–1624 (2018).

7. Capper, D. et al. DNA methylation-based classification of central nervous system tumours. Nature 555, 469–474 (2018).

8. Sturm, D. et al. Hotspot mutations in H3F3A and IDH1 define distinct epigenetic and biological subgroups of glioblastoma. Cancer Cell 22, 425–437 (2012).

9. Kim, T.-M., Huang, W., Park, R., Park, P. J. & Johnson, M. D. A developmental taxonomy of glioblastoma defined and maintained by MicroRNAs. Cancer Res. 71, 3387–3399 (2011).

10. Ciafrè, S. A. et al. Extensive modulation of a set of microRNAs in primary glioblastoma. Biochem. Biophys. Res. Commun. 334, 1351–1358 (2005).

11. Kouzarides, T. Chromatin modifications and their function. Cell 128, 693–705 (2007).

12. Bannister, A. J. & Kouzarides, T. Regulation of chromatin by histone modifications. Cell Res. 21, 381–395 (2011).

13. Tessarz, P. & Kouzarides, T. Histone core modifications regulating nucleosome structure and dynamics. Nat. Rev. Mol. Cell Biol. 15, 703–708 (2014).

14. Bonn, S. et al. Tissue-specific analysis of chromatin state identifies temporal signatures of enhancer activity during embryonic development. Nat. Genet. 44, 148–156 (2012).

15. Bogdanovic, O. et al. Dynamics of enhancer chromatin signatures mark the transition from pluripotency to cell specification during embryogenesis. Genome Res. 22, 2043–2053 (2012).

16. Bernstein, B. E. et al. A bivalent chromatin structure marks key developmental genes in embryonic stem cells. Cell 125, 315–326 (2006).

17. Xu, L. et al. Topography of transcriptionally active chromatin in glioblastoma. Sci. Adv. 7, eabd4676 (2021).

18. Liau, B. B. et al. Adaptive Chromatin Remodeling Drives Glioblastoma Stem Cell Plasticity and Drug Tolerance. Cell Stem Cell 20, 233-246.e7 (2017).

19. Lin, B. et al. Global analysis of H3K4me3 and H3K27me3 profiles in glioblastoma stem cells and identification of SLC17A7 as a bivalent tumor suppressor gene. Oncotarget 6, 5369–5381 (2015).

20. Hall, A. W. et al. Bivalent Chromatin Domains in Glioblastoma Reveal a Subtype-Specific Signature of Glioma Stem Cells. Cancer Res. 78, 2463–2474 (2018).

21. Stępniak, K. et al. Mapping chromatin accessibility and active regulatory elements reveals pathological mechanisms in human gliomas. Nat. Commun. 12, 3621 (2021).

22. Mallm, J.-P., et al. Glioblastoma initiating cells are sensitive to histone demethylase inhibition due to epigenetic deregulation. Int. J. Cancer 146, 1281–1292 (2020).

23. Vinel, C. et al. Comparative epigenetic analysis of tumour initiating cells and syngeneic EPSC-derived neural stem cells in glioblastoma. Nat. Commun. 12, 6130 (2021).

24. Miller, J. L. & Grant, P. A. The Role of DNA Methylation and Histone Modifications in Transcriptional Regulation in Humans. Subcell. Biochem. 61, 289–317 (2013).

25. Wu, K. et al. Dynamics of histone acetylation during human early embryogenesis. Cell Discov. 9, 1–24 (2023).

26. Garciaz, S. et al. Epigenetic down-regulation of the HIST1 locus predicts better prognosis in acute myeloid leukemia with NPM1 mutation. Clin. Epigenetics 11, 141 (2019).

27. Giebel, N. et al. USP42 protects ZNRF3/RNF43 from R-spondin-dependent clearance and inhibits Wnt signalling. EMBO Rep. 22, e51415 (2021).

28. J, S., et al. Amplification and differential expression of members of the erbB-gene family in human glioblastoma. J. Neurooncol. 22, (1994).

29. Logan, C. Y. & Nusse, R. The Wnt signaling pathway in development and disease. Annu. Rev. Cell Dev. Biol. 20, 781–810 (2004).

30. Byrne, K. F., Pal, A., Curtin, J. F., Stephens, J. C. & Kinsella, G. K. G-protein-coupled receptors as therapeutic targets for glioblastoma. Drug Discov. Today 26, 2858–2870 (2021).

31. Griffin, M., Khan, R., Basu, S. & Smith, S. Ion Channels as Therapeutic Targets in High Grade Gliomas. Cancers 12, E3068 (2020).

32. Elsers, D., Temerik, D. F., Attia, A. M., Hadia, A. & Hussien, M. T. Prognostic role of ALK-1 and h-TERT expression in glioblastoma multiforme: correlation with ALK gene alterations. J. Pathol. Transl. Med. 55, 212–224 (2021).

33. Dong, Z. et al. Targeting Glioblastoma Stem Cells through Disruption of the Circadian Clock. Cancer Discov. 9, 1556 (2019).

34. Dong, Z. et al. Targeting Glioblastoma Stem Cells through Disruption of the Circadian Clock. Cancer Discov. 9, 1556–1573 (2019).

35. Pang, L. et al. Circadian regulator CLOCK promotes tumor angiogenesis in glioblastoma. Cell Rep. 42, 112127 (2023).

36. Chen, P. et al. Circadian Regulator CLOCK Recruits Immune-Suppressive Microglia into the GBM Tumor Microenvironment. Cancer Discov. 10, 371–381 (2020).

37. Khan, S. et al. Intrinsic Interferon Signaling Regulates the Cell Death and Mesenchymal Phenotype of Glioblastoma Stem Cells. Cancers 13, 5284 (2021).

38. Kang, J. et al. Ribosomal proteins and human diseases: molecular mechanisms and targeted therapy. Signal Transduct. Target. Ther. 6, 323 (2021).

39. Ernst, J. & Kellis, M. Chromatin-state discovery and genome annotation with ChromHMM. Nat.Protoc. 12, 2478–2492 (2017).

40. Gopi, L. K. & Kidder, B. L. Integrative pan cancer analysis reveals epigenomic variation in cancer type and cell specific chromatin domains. Nat. Commun. 12, 1419 (2021).

41. Hattori, E. Y. et al. A RUNX-targeted gene switch-off approach modulates the BIRC5/PIF1-p21 pathway and reduces glioblastoma growth in mice. *Commun*. Biol. 5, 939 (2022).

42. Wang, H. et al. NF-κB induces miR-148a to sustain TGF-β/Smad signaling activation in glioblastoma. Mol. Cancer 14, 2 (2015).

43. Haynes, H. R. et al. The transcription factor PPARα is overexpressed and is associated with a favourable prognosis in IDH-wildtype primary glioblastoma. Histopathology 70, 1030–1043 (2017).

44. Hua, W. et al. Suppression of glioblastoma by targeting the overactivated protein neddylation pathway. Neuro-Oncol. 17, 1333–1343 (2015).

45. Liang, Q. et al. Inhibition of basigin expression in glioblastoma cell line via antisense RNA reduces tumor cell invasion and angiogenesis. Cancer Biol. Ther. 4, 759–762 (2005).

46. Drucker, K. L., Gianinni, C., Decker, P. A., Diamandis, E. P. & Scarisbrick, I. A. Prognostic significance of multiple kallikreins in high-grade astrocytoma. BMC Cancer 15, 565 (2015).

47. Bhargava, S., Patil, V., Mahalingam, K. & Somasundaram, K. Elucidation of the genetic and epigenetic landscape alterations in RNA binding proteins in glioblastoma. Oncotarget 8, 16650– 16668 (2017).

48. Etcheverry, A. et al. DNA methylation in glioblastoma: impact on gene expression and clinical outcome. BMC Genomics 11, 701 (2010).

49. Zhang, Y. et al. MET Inhibition Elicits PGC1α-Dependent Metabolic Reprogramming in Glioblastoma. Cancer Res. 80, 30–43 (2020).

50. Lim, E.-J., Suh, Y., Kim, S., Kang, S.-G. & Lee, S.-J. Force-mediated proinvasive matrix remodeling driven by tumor-associated mesenchymal stem-like cells in glioblastoma. BMB Rep. 51, 182–187 (2018).

51. Mottaghitalab, F., Lanjanian, H. & Masoudi-Nejad, A. Revealing transcriptional and post-transcriptional regulatory mechanisms of γ-glutamyl transferase and keratin isoforms as novel cooperative biomarkers in low-grade glioma and glioblastoma multiforme. Genomics 113, 2623– 2633 (2021).

52. DeBruyne, J. P., Weaver, D. R. & Reppert, S. M. CLOCK and NPAS2 have overlapping roles in the suprachiasmatic circadian clock. Nat. Neurosci. 10, 543–545 (2007).

53. Cotto, K. C. et al. DGIdb 3.0: a redesign and expansion of the drug-gene interaction database. Nucleic Acids Res. 46, D1068–D1073 (2018).

54. Albright, A. L., Marton, L. J., Lubich, W. P. & Reigel, D. H. CSF polyamines in childhood. Arch. Neurol. 40, 237–240 (1983).

55. Duranton, B. et al. Cytotoxic effects of the polyamine oxidase inactivator MDL 72527 to two human colon carcinoma cell lines SW480 and SW620. Cell Biol. Toxicol. 18, 381–396 (2002).

56. Zhang, D. et al. GABAergic signaling facilitates breast cancer metastasis by promoting ERK1/2-dependent phosphorylation. Cancer Lett. 348, 100–108 (2014).

57. Mack, S. C. et al. Chromatin landscapes reveal developmentally encoded transcriptional states that define human glioblastoma. J. Exp. Med. 216, 1071–1090 (2019).

58. Ye, Y. et al. Long Noncoding RNA CCAL Promotes Papillary Thyroid Cancer Progression by Activation of NOTCH1 Pathway. Oncol. Res. 26, 1383–1390 (2018).

59. Bao, S. et al. Glioma stem cells promote radioresistance by preferential activation of the DNA damage response. Nature 444, 756–760 (2006).

60. Neftel, C. et al. An Integrative Model of Cellular States, Plasticity, and Genetics for Glioblastoma. Cell 178, 835-849.e21 (2019).

61. Elhamamsy, A. R., Metge, B. J., Alsheikh, H. A., Shevde, L. A. & Samant, R. S. Ribosome Biogenesis: A Central Player in Cancer Metastasis and Therapeutic Resistance. Cancer Res. 82, 2344–2353 (2022).

62. Dulken, B. W., Leeman, D. S., Boutet, S. C., Hebestreit, K. & Brunet, A. Single-Cell Transcriptomic Analysis Defines Heterogeneity and Transcriptional Dynamics in the Adult Neural Stem Cell Lineage. Cell Rep. 18, 777–790 (2017).

63. De Boeck, A. et al. Glioma-derived IL-33 orchestrates an inflammatory brain tumor microenvironment that accelerates glioma progression. Nat. Commun. 11, 4997 (2020).

64. Gangoso, E. et al. Glioblastomas acquire myeloid-affiliated transcriptional programs via epigenetic immunoediting to elicit immune evasion. Cell 184, 2454-2470.e26 (2021).

65. Curry, R. N. & Glasgow, S. M. The Role of Neurodevelopmental Pathways in Brain Tumors. Front. Cell Dev. Biol. 9, 659055 (2021).

66. Liu, G. et al. Analysis of gene expression and chemoresistance of CD133+ cancer stem cells in glioblastoma. Mol. Cancer 5, 67 (2006).

67. Gong, Y. et al. Insulin-mediated signaling promotes proliferation and survival of glioblastoma through Akt activation. Neuro-Oncol. 18, 48–57 (2016).

68. Noch, E. K. et al. Insulin feedback is a targetable resistance mechanism of PI3K inhibition in glioblastoma. Neuro-Oncol. 25, 2165–2176 (2023).

69. Hartwell, K. A. et al. The Spemann organizer gene, Goosecoid, promotes tumor metastasis. Proc. Natl. Acad. Sci. U. S. A. 103, 18969–18974 (2006).

70. De Angelis, M. L. et al. An organoid model of colorectal circulating tumor cells with stem cell features, hybrid EMT state and distinctive therapy response profile. J. Exp. Clin. Cancer Res. CR 41, 86 (2022).

71. Xue, T.-C., et al. Goosecoid Promotes the Metastasis of Hepatocellular Carcinoma by Modulating the Epithelial-Mesenchymal Transition. PLOS ONE 9, e109695 (2014).

72. Aiello, N. M. & Stanger, B. Z. Echoes of the embryo: using the developmental biology toolkit to study cancer. Dis. Model. Mech. 9, 105–114 (2016).

73. Pollard, S. M. et al. Glioma stem cell lines expanded in adherent culture have tumor-specific phenotypes and are suitable for chemical and genetic screens. Cell Stem Cell 4, 568–580 (2009).

74. Yang, J. et al. Establishment of mouse expanded potential stem cells. Nature 550, 393–397 (2017).

75. Langmead, B. & Salzberg, S. L. Fast gapped-read alignment with Bowtie 2. Nat. Methods 9, 357– 359 (2012).

76. Li, H. et al. The Sequence Alignment/Map format and SAMtools. Bioinforma. Oxf. Engl. 25, 2078–2079 (2009).

77. Ramírez, F. et al. deepTools2: a next generation web server for deep-sequencing data analysis. Nucleic Acids Res. 44, W160–165 (2016).

78. Zhang, Y. et al. Model-based analysis of ChIP-Seq (MACS). Genome Biol. 9, R137 (2008).

79. Lawrence, M. et al. Software for computing and annotating genomic ranges. PLoS Comput. Biol. 9, e1003118 (2013).

80. Yu, G., Wang, L.-G. & He, Q.-Y. ChIPseeker: an R/Bioconductor package for ChIP peak annotation, comparison and visualization. Bioinforma. Oxf. Engl. 31, 2382–2383 (2015).

81. Ocampo, A. et al. In Vivo Amelioration of Age-Associated Hallmarks by Partial Reprogramming. Cell 167, 1719-1733.e12 (2016).

82. Ernst, J. & Kellis, M. ChromHMM: automating chromatin-state discovery and characterization. Nat. Methods 9, 215–216 (2012).

83. Wang, Q., et al. Exploring Epigenomic Datasets by ChIPseeker. Curr. Protoc. 2, e585 (2022).

84. Raudvere, U. et al. g:Profiler: a web server for functional enrichment analysis and conversions of gene lists (2019 update). Nucleic Acids Res. 47, W191–W198 (2019).

